# Novel combination of CRISPR-based gene drives eliminates resistance and localises spread

**DOI:** 10.1101/2020.08.27.266155

**Authors:** Nicky R. Faber, Gus R. McFarlane, R. Chris Gaynor, Ivan Pocrnic, C. Bruce A. Whitelaw, Gregor Gorjanc

**Author notes:** **For correspondence:** (NRF). **Present address:** Highlander Lab, Easter Bush Campus, Midlothian, EH25 9RG, United Kingdom (Nicky R. Faber, Ivan Pocrnic, Gregor Gorjanc). Laboratory of Genetics, Droevendaalsesteeg 4, 6708PB Wageningen, The Netherlands (Nicky R. Faber). Whitelaw Lab, Easter Bush Campus, Midlothian, EH25 9RG, United Kingdom. Alphagenes Group, Easter Bush Campus, Midlothian, EH25 9RG, United Kingdom (R. Chris Gaynor).

## Abstract

Invasive species are among the major driving forces behind biodiversity loss. Gene drive technology may offer a humane, efficient and cost-effective method of control. For safe and effective deployment it is vital that a gene drive is both self-limiting and can overcome evolutionary resistance. We present HD-ClvR, a novel combination of CRISPR-based gene drives that eliminates resistance and localises spread. As a case study, we model HD-ClvR in the grey squirrel (*Sciurus carolinensis*), which is an invasive pest in the UK and responsible for both biodiversity and economic losses. HD-ClvR combats resistance allele formation by combining a homing gene drive with a cleave-and-rescue gene drive. The inclusion of a self-limiting daisyfield gene drive allows for controllable localisation based on animal supplementation. We use both randomly mating and spatial models to simulate this strategy. Our findings show that HD-ClvR can effectively control a targeted grey squirrel population, with little risk to other populations. HD-ClvR offers an efficient, self-limiting and controllable gene drive for managing invasive pests.

## Introduction

CRISPR-based gene drives have the potential to address problems in public health, agriculture and conservation, including the control of invasive species (***Esvelt et al., 2014***). Invasive species impact livelihoods, have severe economic consequences, and are among the major driving forces behind biodiversity loss (***Mooney, 2005***; ***Pejchar and Mooney, 2009***; ***Sala et al., 2000***). Current control methods such as shooting, trapping, and poisoning are labour-intensive, inhumane, expensive, and ineffective in dealing with the scope of the problem in most situations (***Luque et al., 2014***; ***Campbell et al., 2015***; ***Gurnell and Pepper, 2016***). Examples of damaging invasive species as a result of human mediated introduction include rabbits and cane toads in Australia, Asian carp in the US, and the grey squirrel and American mink in the UK.

In this study, we use the grey squirrel (*Sciurus carolinensis*) that is considered invasive in the UK as a case study for gene drive population control. First introduced in the 19^th^ century, the grey squirrel is now widely distributed across the UK (***Middleton, 1930***). Since their introduction there has been a major decline in native red squirrels (*Sciurus vulgaris*). Grey squirrels are both larger and more aggressive than red squirrels and are passive carriers of Squirrelpox virus, which is lethal to red squirrels (***Tompkins et al., 2002***). Without intervention, red squirrels could be lost from the UK mainland within the next few decades (***England, 2010***). In addition to their impact on native red squirrels, grey squirrels also suppress natural forest regeneration through bark stripping of trees (***Mountford et al., 1999***) and likely have a negative impact on biodiversity of native woodland birds by preying on eggs and chicks (***Hewson and Fuller, 2003***). As an invasive pest they are estimated to cost the UK economy more than £14 million per year by debarking trees, gnawing through electricity cables and other forms of property damage (***Williams et al., 2010***). A manageable and robust grey squirrel control strategy remains to be established (***Gurnell and Pepper, 2016***).

CRISPR-based gene drives may offer a humane, efficient, species-specific and cost-effective method for controlling invasive species, including grey squirrels in the UK (***Prowse et al., 2017***; ***McFarlane et al., 2018***); filling a distinct void in the conservation toolbox. Broadly, a gene drive skews the inheritance ratio of an allele towards a super-Mendelian rate and therefore drives itself to spread quickly through a population (***Burt, 2003***). The CRISPR-Cas system that these gene drives are based on comprises two components: a guide RNA (gRNA) and a nonspecific Cas nuclease (***Cong et al., 2013***). The gRNA directs the Cas nuclease to a specific sequence in the genome where it generates a double stranded break. Several synthetic CRISPR-based gene drives have been proposed with three major types aimed at population control: homing, X-shredder and cleave-and-rescue (***Figure 1***) (***Champer et al., 2016***). A homing gene drive works through a process called ‘homing’ (***Esvelt et al., 2014***). The system utilises germline-specific expression of CRISPR-Cas and subsequent cleavage in the germline, which leads to homology-directed repair (HDR) copying the gene drive element onto the homologous chromosome. By locating the homing gene drive cassette within the coding sequence of a haplosufficient female fertility gene, thereby disrupting the gene’s function, female somatic homozygotes will be infertile. As population growth is typically controlled by female reproductive performance (***Burt, 2003***), the population will decline in size due to an increasing number of infertile females within the population. X-shredder gene drive specifically expresses CRISPR-Cas from the Y-chromosome during spermatogenesis to shred the X-chromosome at multiple locations beyond repair (***Galizi et al., 2016***). Therefore, only Y-bearing sperm mature and all or most offspring of an X-shredder father will inherit a gene drive harbouring Y-chromosome and be male. This eventually leads to a population decline due to the lack of breeding females. Cleave-and-rescue gene drive uses CRISPR-Cas to cleave an essential gene while also supplying a recoded, uncleavable ‘rescue’ copy of this gene within the gene drive cassette (***Oberhofer et al., 2019***). Therefore, offspring must inherit the gene drive to be viable. This system can be used to disrupt the function of a female fertility gene.

**Figure 1.**
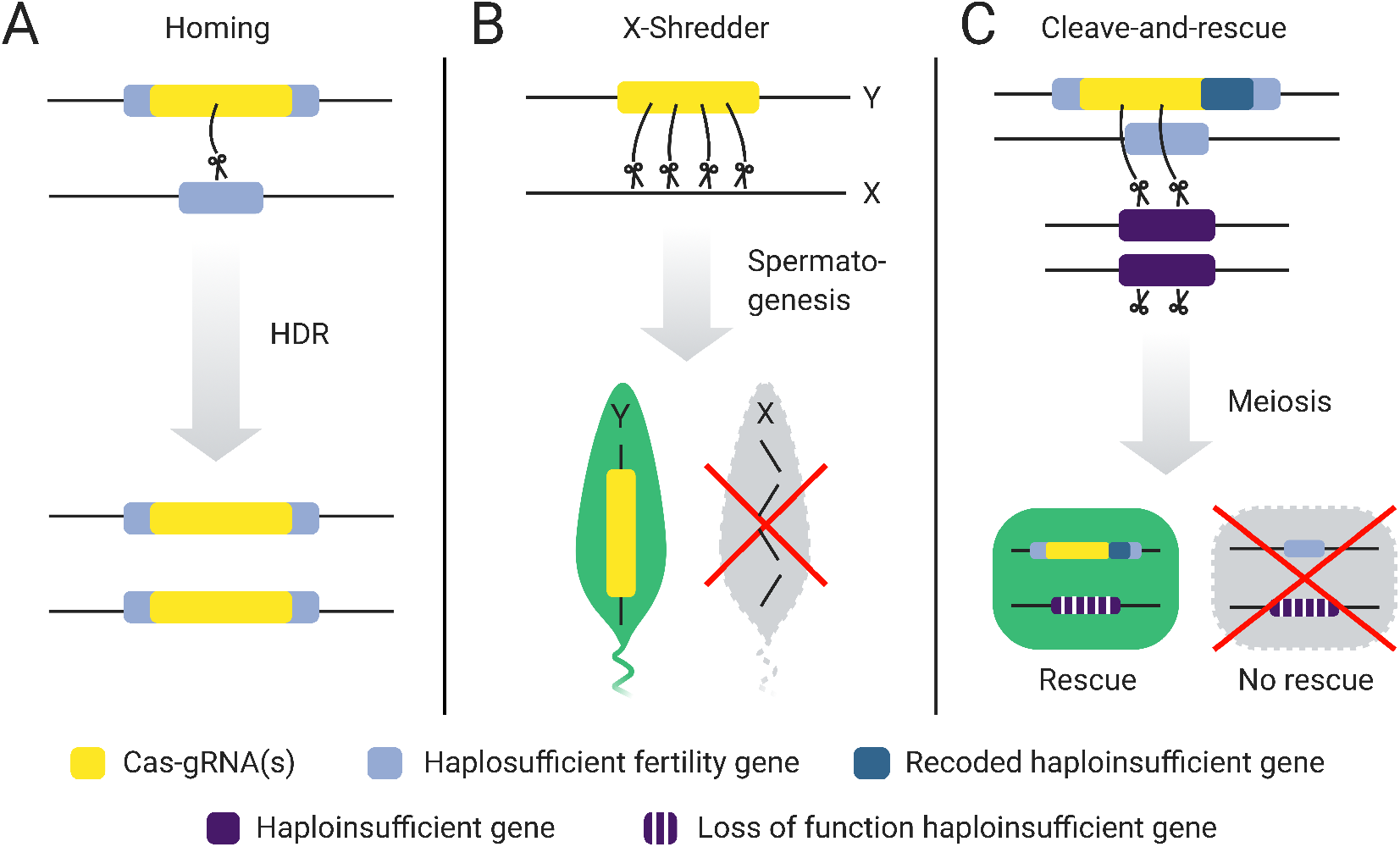
Three CRISPR-based gene drives for population suppression. **A)** Homing. A homing gene drive works by copying itself onto the homologous chromosome in the germline by directing Cas-gRNA(s) to cut a target site, which is repaired via homology directed repair (HDR). Therefore, all or most offspring inherit the gene drive. By locating Cas-gRNA(s) in the coding sequence of a haplosufficient female fertility gene, a female is fertile in homozygous state. All females are infertile once the gene drive allele is fixed leading to suppression of the population. **B)** X-shredder. During spermatogenesis, Cas-gRNA(s) are expressed from the Y-chromosome and shred the X-chromosome beyond repair. Therefore, all or most offspring from an X-shredder father will be shredder males. Population suppression is achieved by skewing the sex-ratio in favour of males. **C)** Cleave-and-rescue. In the germline, Cas-gRNA(s) breaks an essential haploinsufficient gene whilst also supplying a recoded rescue version of this gene in the gene drive cassette. Therefore, only offspring which inherit the rescue within the gene drive are viable. Like the homing gene drive, the cleave-and-rescue gene drive can be located inside a haplosufficient female fertility gene, thereby making somatic homozygote females infertile and achieving population suppression.

Although all three population suppression gene drives are elegant and promising, they all face technical challenges. Homing gene drives face two major challenges. First, during *in vivo* testing, the formation of resistance alleles which block homing have been observed (***Unckless et al., 2017***; ***Champer et al., 2017***). Resistance alleles can form through non-homologous end joining (NHEJ) instead of the desired homology-directed repair during homing. A potential solution is gRNA multiplexing (***Prowse et al., 2017***), but this is likely to reduce homing efficiency (***Champer et al., 2018***, ***2020b***). Second, a homing gene drive that was not hindered by resistant alleles could theoretically spread indefinitely, thereby compromising global ecosystem safety. To address this concern, approaches to make gene drives self-limiting have been divised, including versions called ‘daisy drives’ (***Esvelt and Gemmell, 2017***; ***Noble et al., 2019***; ***Min et al., 2017a***,b). Most daisy drives are complex and likely difficult to engineer, however, the ‘daisyfield’ daisy drive is an exception and provides a straightforward mechanism to limit spread. In a daisyfield gene drive, the gRNAs are scattered throughout the genome (forming a daisyfield) (***Min et al., 2017b***). These daisy elements are inherited in a Mendelian fashion, and therefore, offspring inherits half of the daisy elements from each parent. Thus, the gene drive stops spreading as the daisyfield is diluted through matings with wildtype individuals. Once all daisy elements have disappeared, all elements of the gene drive will likely also disappear due to negative selection drift (as homozygotes are infertile). This is desirable in case gene drive individuals spread to a non-target population. In a population where further spread is required, gene drive individuals with a complete daisyfield can be supplemented to keep the gene drive spreading. The rate and extent of suppression can be controlled by the number of gene drive animals supplemented and how many daisy elements the introduced animals carry. In contrast to homing gene drive, X-shredder gene drives face problems with the formation of a population equilibrium, depending on shredding efficiency (***Beaghton et al., 2017***; ***Champer et al., 2019***). Furthermore, a major challenge in developing X-shredder in mammals is the identification of a highly-specific spermatogenesis promoter to drive Cas-gRNA expression (***McFarlane et al., 2018***). Cleave-and-rescue gene drives have the advantage that multiplexing does not reduce efficiency as there is no homing involved, and therefore, the formation of resistance alleles is limited. Furthermore, cleave-and-rescue gene drives also show density-dependent dynamics, which can be exploited to keep the gene drive contained (***Champer et al., 2020a***). This poses practical challenges as it requires an accurate estimate of population size and the release of a large number of animals simultaneously.

A population control gene drive system that is effective, self-limiting, and controllable has yet to be designed. In this study, we present HD-ClvR, a novel combination of gene drives that eliminates resistance, is self-limiting, and can be controlled in a reliable manner. HD-ClvR is composed of homing (H), daisyfield (D), and cleave-and-rescue (ClvR) gene drives. Our modelling in grey squirrel demonstrates the strategy is highly efficient and overcomes the ongoing issue of resistance allele formation of homing gene drives. The daisyfield gene drive ensures self-limitation and allows for controlled, localised spread. Therefore, HD-ClvR could effectively control a targeted grey squirrel population, with little risk to other populations. Our analysis includes a randomly mating population and a spatially distributed population, which mimics the UK grey squirrel, though it can be adapted to other species. This study provides the first promising steps towards the development and testing of HD-ClvR.

## Results

HD-ClvR is a combination of three gene drives: homing, daisyfield, and cleave-and-rescue. Our randomly mating and spatial modelling of this strategy in grey squirrel illustrates that HD-ClvR can effectively eliminate resistance allele formation, allows for optimised gRNA multiplexing, improves efficiency over standard cleave-and-rescue drives, and is both self-limiting and controllable. We find that the placement of supplemented animals significantly impacts the effectiveness of HD-ClvR, but that this is not prohibitive to the spread of the gene drive and that an effective placement strategy can achieve a rate of gene drive spread close to a randomly mating population.

### Eliminating resistance alleles

By combining a homing gene drive with a cleave-and-rescue gene drive, HD-ClvR eliminates resistance alleles which occasionally form during gene drive homing (***Figure 2A***). This works as follows: as germline homing occurs, both copies of a haploinsufficient essential gene are cleaved, and their function is destroyed through erroneous NHEJ-based repair. However, the homing construct contains a recoded, uncleavable copy of this haploinsufficient gene as a ‘rescue’. For offspring to be viable, they must inherit the gene drive with the rescue to have sufficient expression of the haploinsufficient gene. Offspring that inherit a resistance allele instead of the gene drive will not develop as they lack the rescue gene to compensate for their broken copy of the haploinsufficient gene. This mechanism prevents the spread of resistance alleles.

**Figure 2.**
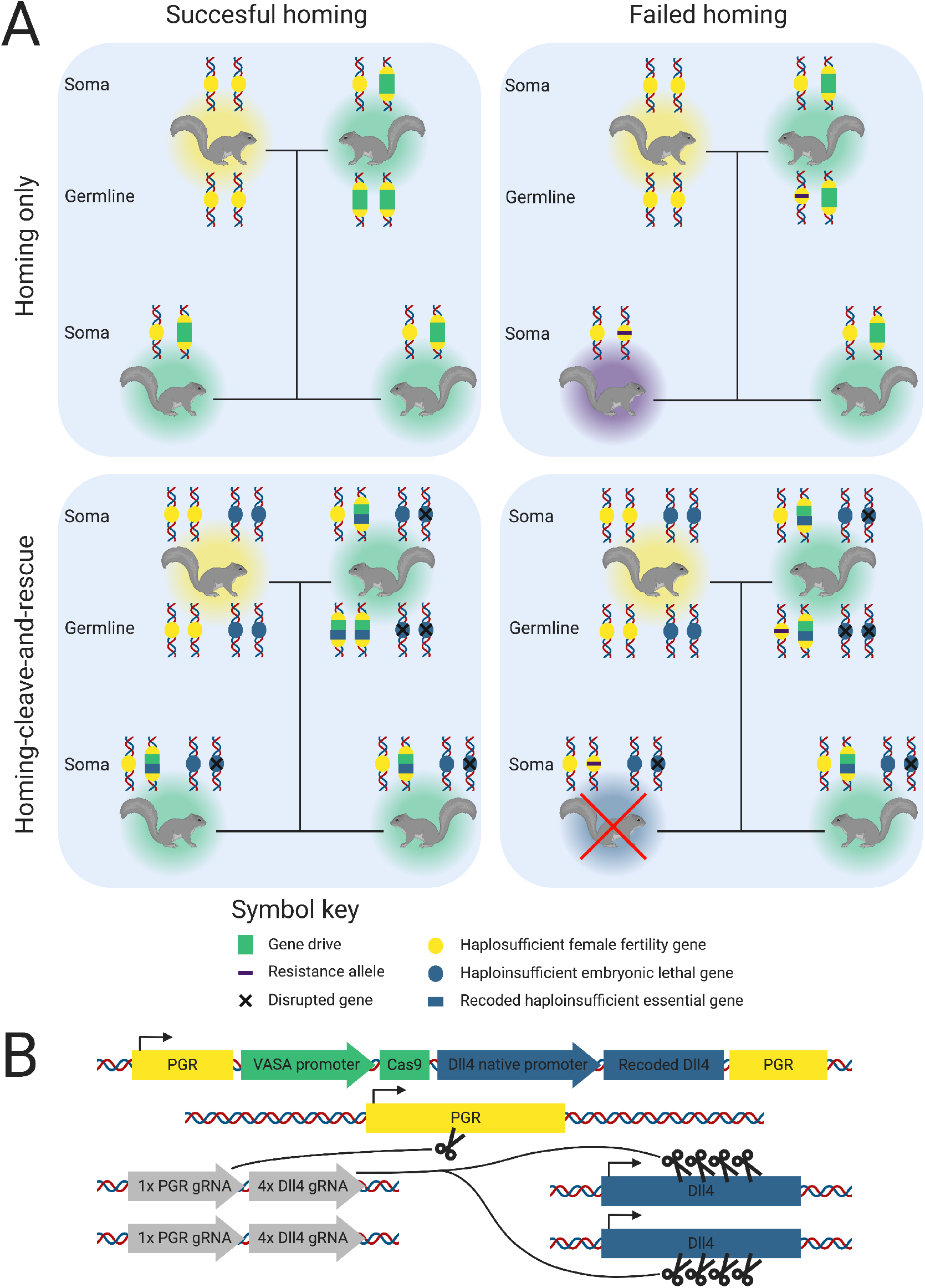
**A)** A comparison of the inheritance scenarios of a homing-only gene drive (top row) and a homing-cleave-and-rescue gene drive (bottom row). The two panels in the left column show inheritance when homing is successful, and the two panels on the right show inheritance when homing fails. Each panel shows two parent squirrels and two offspring, each with the loci relevant for the gene drive. A legend for the gene drive components is provided. Squirrels colour coded halos represent their genotype: yellow = wildtype, turquoise = gene drive, purple = resistant, and blue = non-viable. **B)** A potential HD-ClvR construct for grey squirrel. Colour coding is consistent with **A** and additionally, gRNAs are shown in grey. The gRNAs shown in this figure constitute one daisy element, multiple of these would constitute a daisyfield.

HD-ClvR also allows for independent optimising of gRNA multiplexing for both homing efficiency and resistant allele elimination. Multiplexing gRNAs can overcome resistance allele formation, allowing homing to take place even if some resistant gRNA sites are present. With a standard homing gene drive, the optimal number of gRNAs is a trade-off between homing efficiency and overcoming resistance allele formation. Two gRNAs has been proposed as optimal for homing, with efficiency decreasing when more than two gRNAs are used (***Champer et al., 2020b***). However, to also limit the formation of resistance alleles, the optimal number in the trade-off lies between 4 and 8 (***Champer et al., 2020b***). In contrast, with HD-ClvR it is possible to select the optimal number of gRNAs for homing, while multiplexing several gRNAs within the cleave-and-rescue to reduce the probability of resistance allele formation to effectively zero. Current data suggests four gRNAs is sufficient to prevent resistant allele formation (***Champer et al., 2020b***).

In grey squirrel, we have selected two genes through literature mining which are suitable for HD-ClvR: Progesterone Receptor (PGR) as a haplosufficient female fertility gene and Delta-Like Canonical Notch Ligand 4 (DLL4) as a haploinsufficient essential gene. Both of these genes are conserved across many taxa and could also be used for other invasive species (***Huerta-Cepas et al., 2019***). ***Figure 2B*** shows a candidate HD-ClvR contruct design for grey squirrel control, using 1 gRNA for homing and 4 gRNAs for cleave-and-rescue.

To demonstrate that combining a homing and cleave-and-rescue gene drive can eliminate the formation of resistance alleles, we model a standard homing gene drive, a standard cleave-and-rescue gene drive, and a homing-cleave-and-rescue gene drive in a randomly mating population of grey squirrels over different rates of NHEJ (*P*_*n*_, ***Figure 3***). Like (***Prowse et al., 2017***), we model no fitness cost to heterozygote gene drive animals. Our model uses either 1 or 4 gRNAs to show multiplexing reduces resistance allele formation. For the standard cleave-and-rescue gene drive, we modelled the release of 1000 gene drive squirrels instead of 100 gene drive squirrels, as this form of drive is only effective at a large introduction frequency. A standard homing gene drive was effective at low rates of NHEJ (*P*_*n*_ and 0.1) when multiplexing 4 gRNAs but is inhibited by resistant alleles when only 1 gRNA is used at the same rates of NHEJ. However, at a higher rate of NHEJ (*P*_*n*_ = 0.5), squirrels with resistant alleles rescue the population from standard homing gene drive suppression despite multiplexing 4 gRNAs. In contrast, with a homing-cleave-and-rescue gene drive, resistant alleles are eliminated, and the squirrel population is completely suppressed across all rates of NHEJ when 4 gRNAs are used in the cleave-and-rescue component of the drive. When we compare the three gene drive types in a large population of carrying capacity 30,000 instead of 3,000, we see the same dynamics (***Figure 3–Figure Supplement 1***).

**Figure 3.**
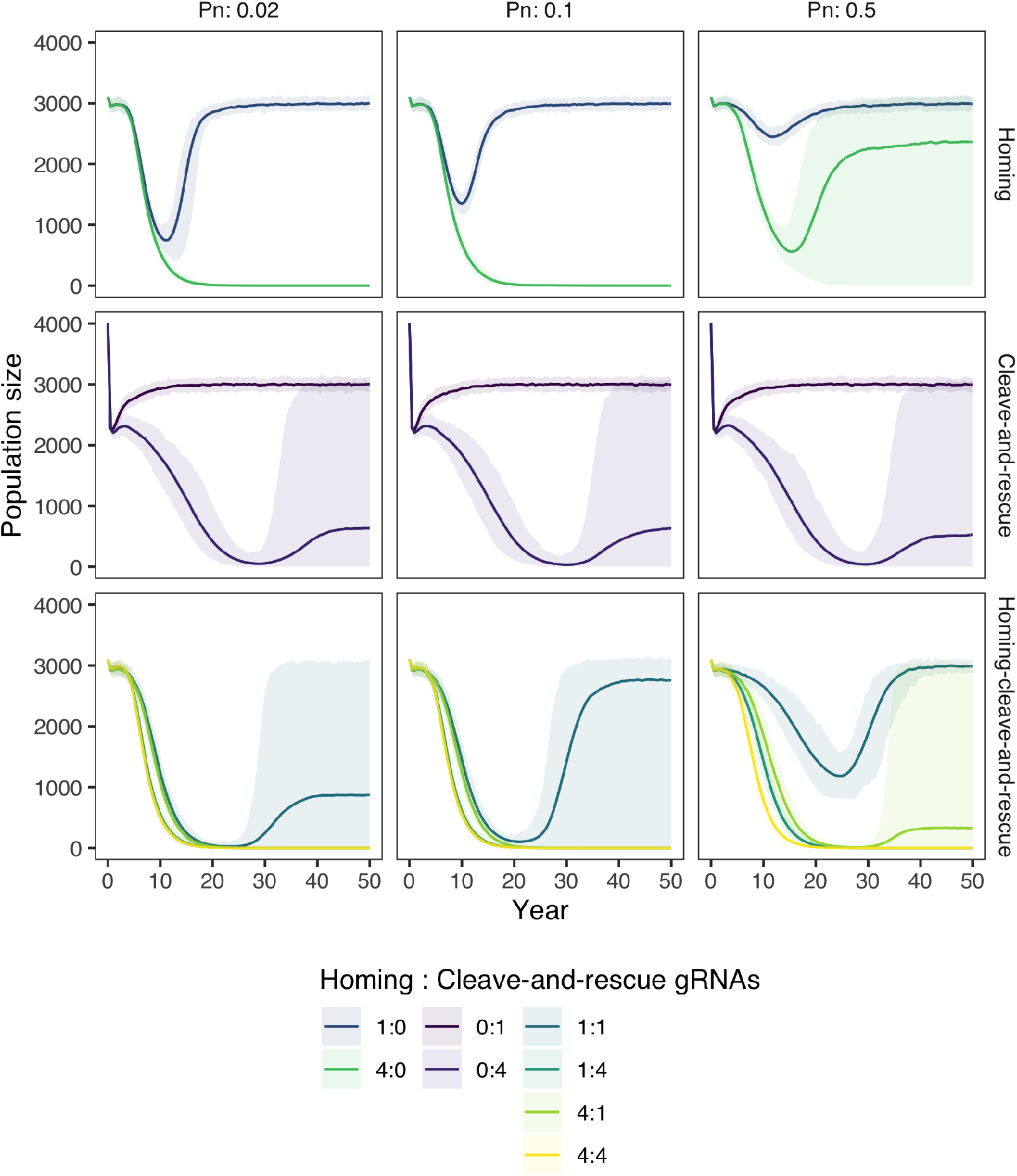
Population size over time after the introduction of gene drive squirrels with either a standard homing, a standard cleave-and-rescue, or a homing-cleave-and-rescue gene drive to a population with carrying capacity 3,000. All simulations are based on a single release of 100 squirrels is done, other than the standard cleave-and-rescue gene drive, which requires a release of 1000 squirrels. Lines represent the average population size over 100 model replications, while opaque ribbons represent the 95% quantiles. The model was run with 3 different rates of NHEJ repair during homing (*Pn*) and with different numbers of gRNAs for the homing and the cleave-and-rescue components of the gene drive. **Figure 3–Figure supplement 1.** The same as ***Figure 3***, but run in a big population with a carrying capacity of 30,000. Introduction numbers were kept at 100, but for the standard cleave-and-rescue gene drives, an introduction frequency of 10,000 was used because of its density dependent mechanics. **Figure 3–Figure supplement 2.** An exploration of which type of gene is best targetted by the cleave-and-rescue part of the homing-cleave-and-rescue gene drive: both-sex infertility or developmental non-viability, and overexpression biologically tolerable or not. Parameters are kept the same as in ***Figure 3***, except that we used 1 gRNA for the homing part of the gene drive, and either 1, 2 or 4 gRNAs for the cleave-and-rescue part.

Although we model the homing gene drive component of HD-ClvR targeting a haplosufficient female fertility gene in this study, HD-ClvR is adaptable and could target any desirable gene to generate a loss of function mutation through insertion disruption or propagate a genetic cargo of interest. The cleave-and-rescue component of the HD-ClvR targets a haploinsufficient developmental gene in this study but this could also be adjusted to a haploinsufficient both-sex infertility gene. Our results suggest it is marginally more efficient to target an embryonic lethal gene (***Figure 3–Figure Supplement 2***), as this prevents infertile resistant individuals from competing with gene drive individuals for resources. From an ethical standpoint the reduction in efficiency when targeting a both-sex fertility gene, instead of an embryonic lethal gene, may be justified by the improved societal and political acceptance for a strategy that evades killing and suppresses through infertility. Additionally, we tested if overexpression of the cleave-and-rescue target gene should be biologically tolerable (***Figure 3–Figure Supplement 2***). We conclude that when multiplexing sufficiently for the cleave-and-rescue part of the gene drive, there is no difference. As can be seen from the dynamics when multiplexing less or not at all, allowing overexpression makes the gene drive initially faster to spread, but also allows resistance alleles to persist in the population.

### Self-limitation and control

A key benefit of HD-ClvR is that by including a daisyfield gene drive, it is self-limiting and can be controlled based on the number of supplemented gene drive animals and number of daisy elements each supplemented animal harbours (***Figure 4***). Unlike a standard homing gene drive, HD-ClvR can control the rate and extent of population suppression and, if required, suppression could be stopped by terminating further animal supplementation. Additionally, HD-ClvR does not require the large initial releases of standard cleave-and-rescue animals, which places pressure on the local ecosystem.

**Figure 4.**
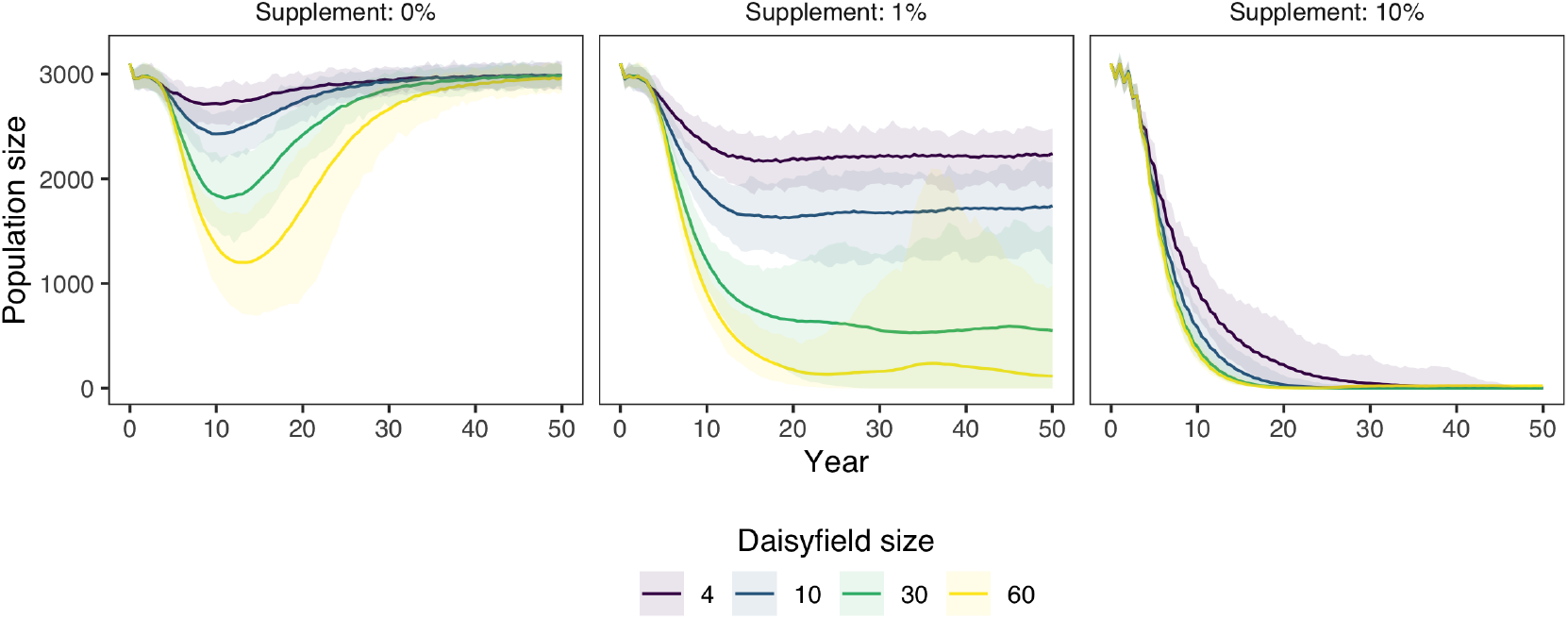
Population size over time after the introduction of 100 squirrels with a HD-ClvR gene drive to a population of carrying capacity 3,000. Lines represent the average population size over 100 model replications, while opaque ribbons represent the 95% quantiles. The model was run with an NHEJ rate (*Pn*) of 0.02, 1 homing gRNA, and 4 cleave-and-rescue gRNAs. Gene drive squirrel supplementation was done yearly, the amount being a percentage (0, 1, or 10%) of the total population size at that moment. **Figure 4–Figure supplement 1.** The same as ***Figure 4***, but run in a big population with a carrying capacity of 30,000. **Figure 4–Figure supplement 2.** The same as ***Figure 4***, but instead of an accurate estimate of the population size for supplementation, a certain level of error is introduced. The error is defined as a normal distribution with the true population size as mean and a certain percentage of the true population size as standard deviation. **Figure 4–Figure supplement 3.** The same as ***Figure 4***, but ran with a range of supplementation amounts and daisyfield sizes. Suppression rate is defined as the proportion of populations (out of the 100 repetitions of the model) that were completely suppressed after 50 years.

Using our randomly mating model, we show in ***Figure 4*** that by including a daisyfield system in a homing-cleave-and-rescue drive to form HD-ClvR, we can efficiently suppress a targeted population, while limiting risk to other populations, especially if those are bigger than the target population (***Figure 4–Figure Supplement 1***). We modelled HD-ClvR with different daisyfield sizes in a population of 3,000 grey squirrel over different rates of annual supplementation following an initial release of 100 HD-ClvR squirrels. The model shows that once the HD-ClvR runs out of daisy elements the population recovers. Therefore, HD-ClvR poses less risk to non-target populations than a standard homing gene drive. With 1% annual supplementation of HD-ClvR squirrels, the population size is reduced and maintained at an equilibrium, and with 10% annual supplementation the targeted population of grey squirrel is removed for all daisyfield sizes. In ***Figure 4–Figure Supplement 2***, we show that it is possible to suppress a population without an accurate estimation of population size, which will be hard to obtain for most wild populations. To find the optimal combination of supplementation rate and daisyfield size, we ran a range of these two parameters and found that 5% supplementation would be sufficient to suppress a population, even with a small daisyfield (***Figure 4–Figure Supplement 3***).

### Spatial dynamics and supplementation of HD-ClvR

To understand the spatial dynamics of homing-cleave-and-rescue drives, initially excluding daisyfield, we modelled this approach in a simple spatial model. Modelling a single release of 100 homing-cleave-and-rescue gene drive squirrels in populations of 3,000 and 30,000 squirrels, the model demonstrated that the spatial life history of grey squirrel allows for the spread of the gene drive (***Figure 5***). We also show that the removal of the target squirrel population is more delayed in the spatial model than in the randomly mating population model. This difference is approximately five years in a small population, and is increased to approximately 15 to 20 years in a big population. To test the sensitivity of our model to two crucial parameters, mating range and migration range, we performed a sensitivity analysis and conclude that the model is sensitive to a decreased mating range, but not to a decreased migration range (***Figure 5–Figure Supplement 1***).

**Figure 5.**
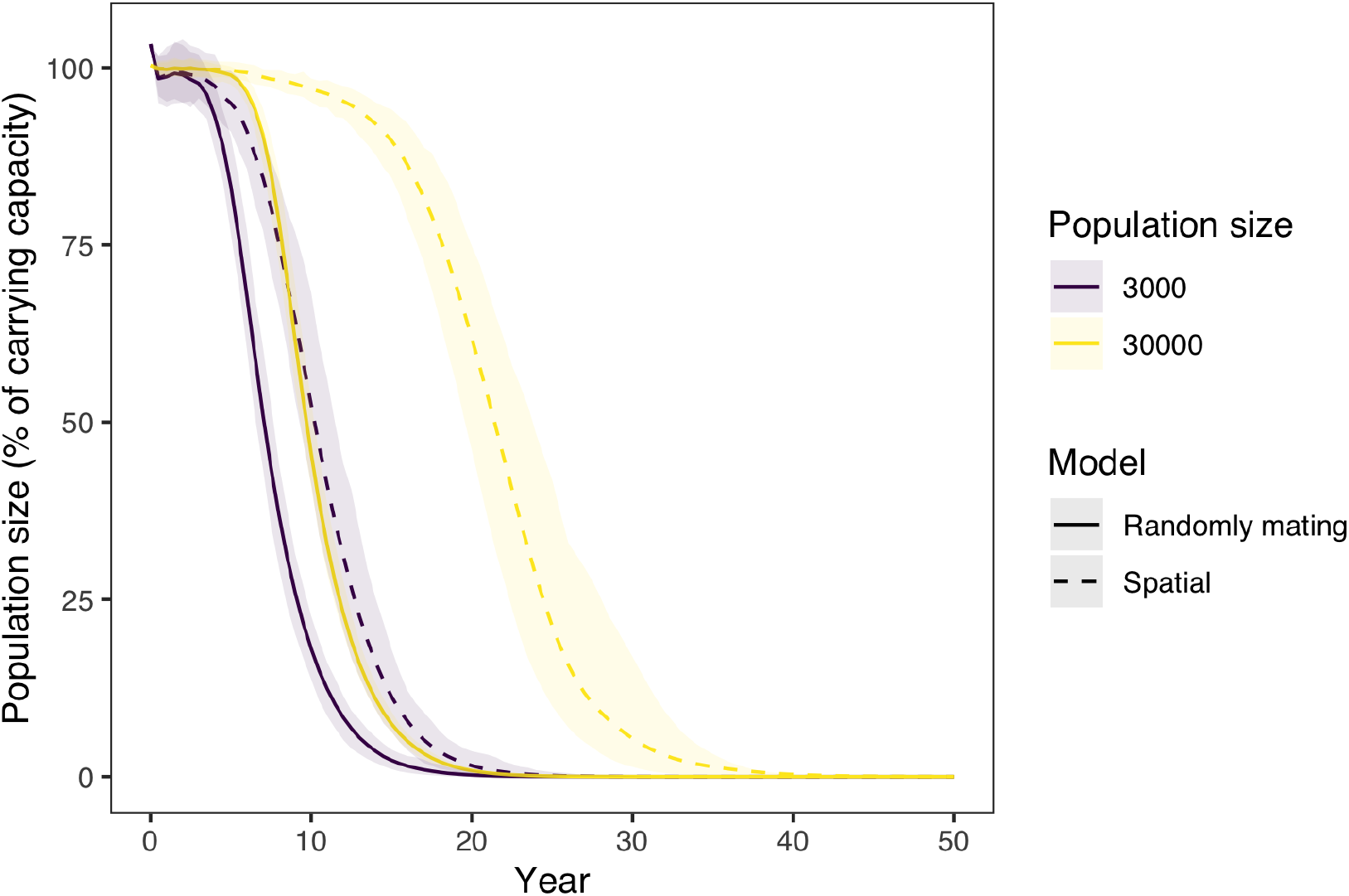
Population size over time after the introduction of 100 squirrels with a homing-cleave-and-rescue gene drive with 1 homing gRNA and 4 cleave-and-rescue gRNAs. The model was run for a randomly mating and a spatial model, and also for a small (carrying capacity 3,000) and large population (carrying capacity 30,000). In the spatial model, gene drive squirrels were placed in the middle of the area. An NHEJ rate (*Pn*) of 0.02 was used. Lines represent the average population size over 100 model replications, while opaque ribbons represent the 95% quantiles. **Figure 5–Figure supplement 1.** The same as the small population with a carrying capacity of 3,000 in a spatial model in ***Figure 5***, but a sensitivity analysis of two crucial parameters: mating range and migration range.

Using our spatial model, we then explored how the placement of supplemented HD-ClvR animals could impact population suppression. We show the impact of different supplementation placement schemes by modelling five strategies: mean of population location, mode of population location, randomly, randomly in 10 groups, and in a moving front (***Figure 6A***). The moving front was implemented such that we start at the bottom and move upwards in ten steps, thereafter, supplementing at the topmost location. As can be seen in ***Figure 6B***, different placement schemes significantly affect the efficiency of the strategy. Placement at the mean population location was least effective and placement of squirrels randomly in 10 groups was most effective. ***Figure 6C*** shows three moments which represent key spatial dynamics of each placement scheme. For animations of the spatial dynamics over the whole timeline, see the animated GIFs (***Figure 6***–***video 1***).

**Figure 6.**
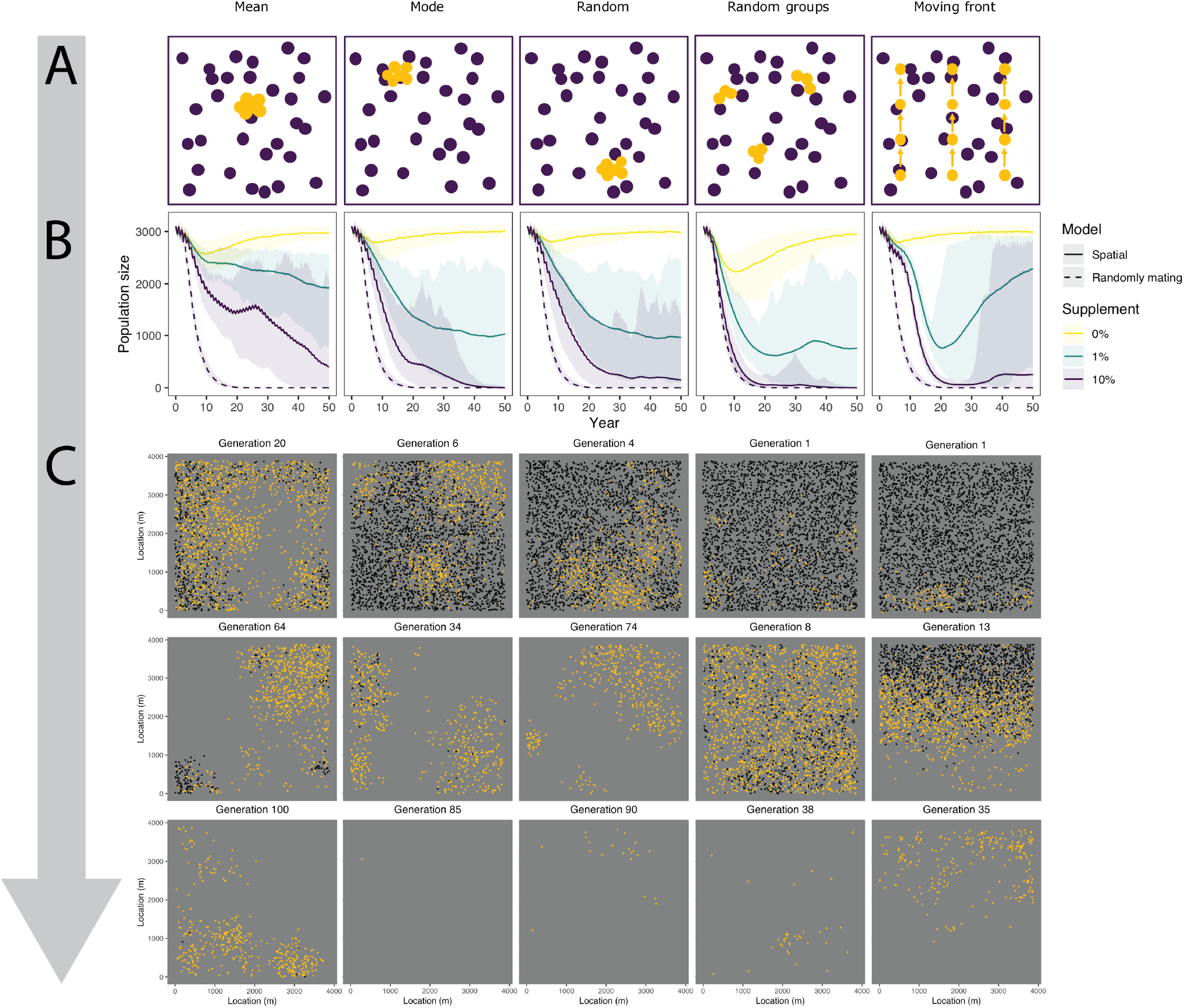
Spatial dynamics of HD-ClvR using different placement schemes. **A)** A schematic overview of the placement schemes. **B)** Population size chemes and amounts of supplementation. We modelled population size over time after the introduction of 100 ive with 1 homing gRNA and 4 cleave-and-rescue gRNAs to a population of carrying capacity 3,000. We modelled aisyfield of size 30. **C)** Three snapshots of moments representing key spatial dynamics at 10% supplementation. See the full animations in ***video 1***. **Figure 6–video 1.** Full animations of the spatial dynamics of HD-ClvR using the five placement schemes (see https://git.ecdf.ed.ac.uk/HighlanderLab_public/nfaber_squirrel_gd/tree/master/Fig6_GIFs). We model the spatial dynamics of a population over time after the introduction of 100 squirrels homing gRNA and 4 cleave-and-rescue gRNAs to a population of carrying capacity 3,000. We modelled an NHEJ ize 30 and a supplementation amount of 10%.

## Discussion

This research presents HD-ClvR, which is a combination of three gene drives: homing, cleave-and-rescue and daisyfield. Our modelling indicates that HD-ClvR overcomes an important trade-off in current homing gene drive designs: the trade-off between resistance allele formation and gene drive efficiency. This strategy benefits from the efficiency of a homing gene drive and the evolutionary stability of cleave-and-rescue gene drive. Due to the inclusion of a daisyfield system, HD-ClvR is self-limiting and can be controlled by supplementation of gene drive animals.

### HD-ClvR compared to other gene drives

Over recent years, many different gene drives have been published and developments have been geared towards both efficiency and safety (***Champer et al., 2016***). An ongoing issue has been the development of resistance alleles. For CRISPR-based homing gene drive there are two fundamental approaches to combat resistance allele formation: careful gRNA targeting and gRNA multiplexing. When a gRNA targets a conserved sequence in a gene, resistance alleles are likely to disrupt gene function through NHEJ repair and will therefore reduce fitness (***Kyrou et al., 2018***). Recently, population suppression was already shown to work with a carefully targeted homing gene drive in contained mosquito populations (***Kyrou et al., 2018***), however, current data suggests that homing might be less efficient in mammals than in insects (***Grunwald et al., 2019***). Very recently, a new preprint has proposed a gene drive very similar to HD-ClvR, which combines a homing and cleave-and-rescue gene drive to combat resistance alleles (***Kandul et al., 2020***).

In addition to targeting conserved sequences, when gRNA multiplexing, resistant allele allele formation is reduced because multiple sites are targeted simultaneously. For homing gene drives, multiplexing has been shown to reduce homing efficiency when more than two gRNAs are used (***Champer et al., 2020b***). In contrast, cleave-and-rescue gene drives do not have this problem, as they do not use homing and can therefore multiplex gRNAs without any efficiency costs. HD-ClvR separates the elimination of resistance alleles and homing efficiency, and therefore gRNAs can be optimised for both goals separately.

To date, most gene drive research has focused on improving the efficiency, however, equally important is the development of strategies that allow for containment, or even reversibility, of the gene drives (***Esvelt and Gemmell, 2017***; ***Marshall and Hay, 2012***). For contained gene drives, density dependence is often used, which requires large numbers of gene drive individuals to be released into a target population to spread (***Edgington and Alphey, 2017***). Therefore, non-target populations are unlikely to be affected by this type of gene drive. However, a large single release of gene drive individuals can put significant pressure on the local ecosystem, and if a population is already at carrying capacity, it may lead to starvation or mass migration of the population. In contrast, HD-ClvR uses ongoing input in the form gene drive animals to control the extent of population suppression and contain spread. Although this comes with increased cost and labour, we believe this is justified by the improved control and safety HD-ClvR could offer over current gene drives.

As stated above, the initial introduction frequency for a standard cleave-and-rescue gene drive in our randomly mating model was increased 10-fold over the other homing-based strategies. This increase is necessary due to the significant cost to the reproduction rate that is incurred when using a standard cleave-and-rescue gene drive. On average, cleave-and-rescue animals will produce 50% less offspring than wild-type animals (***Oberhofer et al., 2019***; ***Champer et al., 2020a***). This significantly slows the spread of the gene drive and due to density dependent dynamics, requires large initial releases of cleave-and-rescue animals for population suppression. With a homing-cleave-and-rescue drive, more offspring inherit the drive and there is less cost to the reproduction rate. Effectively, for homing-cleave-and-rescue, the reproduction rate of gene drive individuals is equal to the homing efficiency (plus half of the homing failure rate, where the gene drive is inherited by chance), which so far has been shown to range from 0.7 to 1 in different organisms (***Kyrou et al., 2018***; ***Gantz et al., 2015***; ***Grunwald et al., 2019***).

### Supplementation

As animal supplementation is a critical component of HD-ClvR, our modelling investigated how daisyfield size and the level and placement of supplemented HD-ClvR animals effects efficiency and safety of population suppression. Optimisation of these parameters can significantly reduce cost and labour, as well as reduce the risk of unwanted impacts on non-target populations. We modelled our supplementation as a percentage of the total population size, therefore the number of individuals needed for supplementation increases linearly with population size. We also want to minimise the risk of non-target populations being impacted by the gene drive, and therefore, there is a trade-off between safety (size of the daisyfield) and cost and labour (level of supplementation required).

The least number of daisy elements that can suppress the population with a realistic level of supplementation, but does not cause any serious issues in non-target populations, should be objectively established through an in-depth risk assessment process. In a larger population however, the spread is slower than in a small one. Therefore, for improved safety and efficiency, gene drives are best applied in small sub-populations separately. The impact of a single introduction, such as a rogue deployment or migration, depends on the population size. The smaller the population, the bigger the impact. This it is a concern when the target population is much larger than the non-target population, but this is not the case for invasive UK grey squirrels and many other invasive species.

The appropriate daisyfield size also depends on the rate of NHEJ (*P*_*n*_) of the gene drive system; the higher the (*P*_*n*_), the more embryonic lethal offspring will arise and the sooner daisyfield burns out. To choose a safe number of daisy elements, we also need an estimate of how many animals a rogue party could obtain, potential breed and add into a non-target population for their own benefit. Overall, each target population and prospective gene drive strategy needs to be considered on a case-by-case basis and include an in-depth multidisciplinary risk assessment process.

When we consider the spatial aspects of a HD-ClvR supplementation programme, the picture becomes more complex. A key factor is the supplementation location of individuals. Obviously, supplementing individuals in a location where the population has already been suppressed will be ineffective. Therefore, different placement strategies can be adopted to keep placing individuals in a relevant area. A monitoring system where not only the size of the population is known, but also the location can significantly help HD-ClvR continue spreading and suppress a targeted population.

In this study, we modelled HD-ClvR using five different supplementation placement strategies in grey squirrel. These were: supplementation at the mean of population location, the mode of population location, randomly, randomly in 10 groups, and in a moving front (***Figure 6A***). With supplementation at the mean of the population location, supplementation started in the middle of the population. After a few generations, a gap appears in the middle due to local suppression. The mean of the populations location still lies in the middle, as can be seen in ***Figure 6C*** at 20 generations. Therefore, supplementation is not effective until the population is also suppressed in another location, thereby shifting the mean. Additionally, when there is a single large patch of the population left and additional smaller clusters, supplementation in the middle of the large patch allows the smaller clusters to recover, as can be seen in ***Figure 6C*** after 64 generations.

With supplementation at the mode of the population location, we supplement in a location where there are many individuals. This placement strategy avoids the problem of supplementing in a location without individuals, either in a doughnut-like spatial population structure or in a multi-patch population. However, this placement strategy still allows small patches to form and recover. Supplementation at a random location theoretically means that supplementation happens uniformly, but in reality, this is not the case. Initially HD-ClvR spreads in multiple locations, but after the population is suppressed in certain regions, supplementation in those regions becomes ineffective. Therefore, at a later stage of population suppression this placement scheme becomes increasingly ineffective.

Supplementation at random locations is more effective when they are broken up into multiple groups (ten in our model). The gene drive spreads in many locations initially like the random single location placement scheme. After significant suppression of the population some but not all of the 10 groups supplemented are at ineffective locations. The groups that are placed at relevant locations are enough to keep the gene drive spreading. In our model supplementation in groups at random locations gets close to the speed at which a gene drive spreads in a non-spatial model.

The moving front placement scheme is very effective initially, as the gene drive spreads uniformly across the front. In this case, supplementation keeps ahead of where the populations is being suppressed. This placement strategy allows the population to recover behind the moving front after effective initial spread and near-complete suppression. To improve efficiency of the moving front strategy, it may be beneficial to include random supplementation behind the moving front to prevent animals from re-establishing.

Finally, in our spatial model, it was evident that there is more uncertainty in levels of population suppression than a randomly mating model leads us to believe. As can be seen in ***Figure 6B***, the 95% quantiles are broader than the quantiles in ***Figure 3***. Therefore, we conclude that to tailor the amount of supplementation, it is vital to closely monitor a population where a gene drive is used.

### Assumptions and future work

Our model works under the following six assumptions. First, our model excludes some complexities of the optimal number of gRNAs for homing. Although our model suggests that multiplexing gRNAs for both the homing and cleave-and-rescue gene drives is most effective, a recent study using a more complex model and *in vivo* data shows that the optimal number of gRNAs to use for homing in *Drosphilia melanogaster* is two. They report a decrease in homing efficiency with more than two gRNAs due to reduced homology and Cas nuclease saturation (***Champer et al., 2020b***). Therefore, our gene drive with four gRNAs for both homing and cleave-and-rescue will likely be less efficient in such a complex model. We suggest using two homing gRNAs and four cleave- and-rescue gRNAs is likely most efficient, while still eliminating all resistance alleles (***Champer et al., 2020b***). It would be prudent to analyse our gene drive in this complex model as well to get a definitive estimate, as Cas saturation is thought to have an influence on gene drive efficiency when multiplexing is used (***Champer et al., 2020b***).

Second, we assumed there was no embryonic Cas-gRNA expression. Embryonic Cas-gRNA expression might be problematic as it leads to resistance allele formation and can interfere with the cleave-and-rescue mechanism by cleaving alleles from the wildtype parent. As our gene drive eliminates resistance alleles, embryonic Cas-gRNA expression may not inhibit spread, depending on the rate. Additionally, if the embryonic Cas-gRNA expression turns out to be more common in grey squirrel or other species, the cleave-and-rescue part of the gene drive can be harnessed with a double rescue mechanism to overcome this issue, as reported by ***Champer et al. (2020a)***.

Third, we did not take other types of resistance alleles into account such as mutations rendering the CRISPR-Cas non-functional. As this is a universal assumption in gene drive research, we will have to await multigenerational studies to see if this is problematic.

Fourth, HD-ClvR has not been tested *in vivo*, which is our next step. The recent preprint on a gene drive very similar to HD-ClvR has performed *in vivo* tests in *Drosophila melanogaster* which showed very efficient conversion rates (***Kandul et al., 2020***). Proof-of-concept testing of HD-ClvR would likely initially occur in *D. melanogaster* and mouse models before progressing to squirrel studies. Also, recent reports have shown that the VASA promoter for Cas expression in homing gene drives is not optimal and further investigation to identify a meiosis-specific germline promoter is needed (***Pfitzner et al., 2020***). Furthermore, non-model species might be difficult to genetically engineer, although grey squirrel embryology will likely follow the extensive knowledge on rodent and farmed animal embryology, and similar reagents and equipment could be used. An important consideration when engineering gene drive is that the modified animals maintain enough wild vigour to survive and breed in a wild population. Promising technologies for generating gene drive harbouring mammals with as little intervention as possible include *in vivo* zygotic delivery of CRISPR reagents by electroporation or viral transduction (***Mehier-Humbert and Guy, 2005***; ***Zhang and Godbey, 2006***).

Fifth, for our spatial modelling, we assumed that an estimation of population size could be made every year, although there is a significant amount of room for error in this estimate. Additionally, for some of our placement schemes, we assumed an accurate estimate of population location. As the random placement in groups scheme turned out most effective, this is not a problem so much as further potential for improvement. Another direction for future spatial work is the modelling of real landscapes, which are more complex than what we modelled in this study (***Bradburd and Ralph, 2019***). In complex landscapes, it might be that gene drive spread is slower or even regionally confined in some situations. Additionally, there might be spatial dynamics to gene drives in general such as ‘chasing’, which is the perpetual escaping and chasing of wildtype and gene drive animals (***Champer et al., 2019***). Further efforts are necessary to create a more realistic spatial model before we can consider using a gene drive.

A final consideration is that the ecological services the grey squirrel and other invasive species provide are largely unchartered. Ecologists need to investigate the ecological services that an invasive species performs and how an abrupt suppression of this invasive population might impact the ecosystem as a whole. We need to consider other restorative measures such as reintroducing native species to fragmented habitats, amongst other ecological interventions (***Rode et al., 2019***). From a regulatory perspective, there is no tested legislative framework for the release of gene drive organisms; and with regard to our test animal it is currently illegal to breed grey squirrels in the UK. Developing these legislative frameworks alongside gene drive research is important. More importantly, the UK needs to continue to broaden public engagement and see whether the public is receptive to the deployment of gene drive technology in parallel to a financial overview of how much it would cost to apply gene drives reflecting our predicted need for supplementation.

### Summary

HD-ClvR offers an efficient, self-limiting, and controllable gene drive strategy. We show that in the spatial model, complete population suppression is achieved approximately five years later than in the randomly mating population model. We then explored how the placement of supplemented animals could impact population suppression. Our results show that spatial dynamics of supplementation placement are not prohibitive to the spread of the gene drive, but that in fact, with an optimised strategy, spread at a rate equal to randomly mating population can be achieved. In our models, we have shown that grey squirrels have a spatial life history which facilitates the spread of a gene drive. Therefore, gene drives could be a valuable tool in the conservation toolbox.

## Methods and Materials

We describe our methods and materials in two sections. The first section details the randomly mating population model, and the second the spatial model. For the modelling, we adopted the work of ***Prowse et al. (2017)*** and implemented new features. This model is an individual-based, stochastic, discrete-time model of a randomly mating population. Per individual, the model keeps track of several characteristics such as age, sex, parents, and the state of genetic loci involved in the gene drive. For each offspring, we model the homing and subsequent inheritance of the gene drive. By running this stochastic model several times, we obtain an impression of the possible outcomes. Several life history parameters of an organism are needed to run this model. The parameters we used to model a grey squirrel population can be seen in ***Table 1***.

**Table 1.**
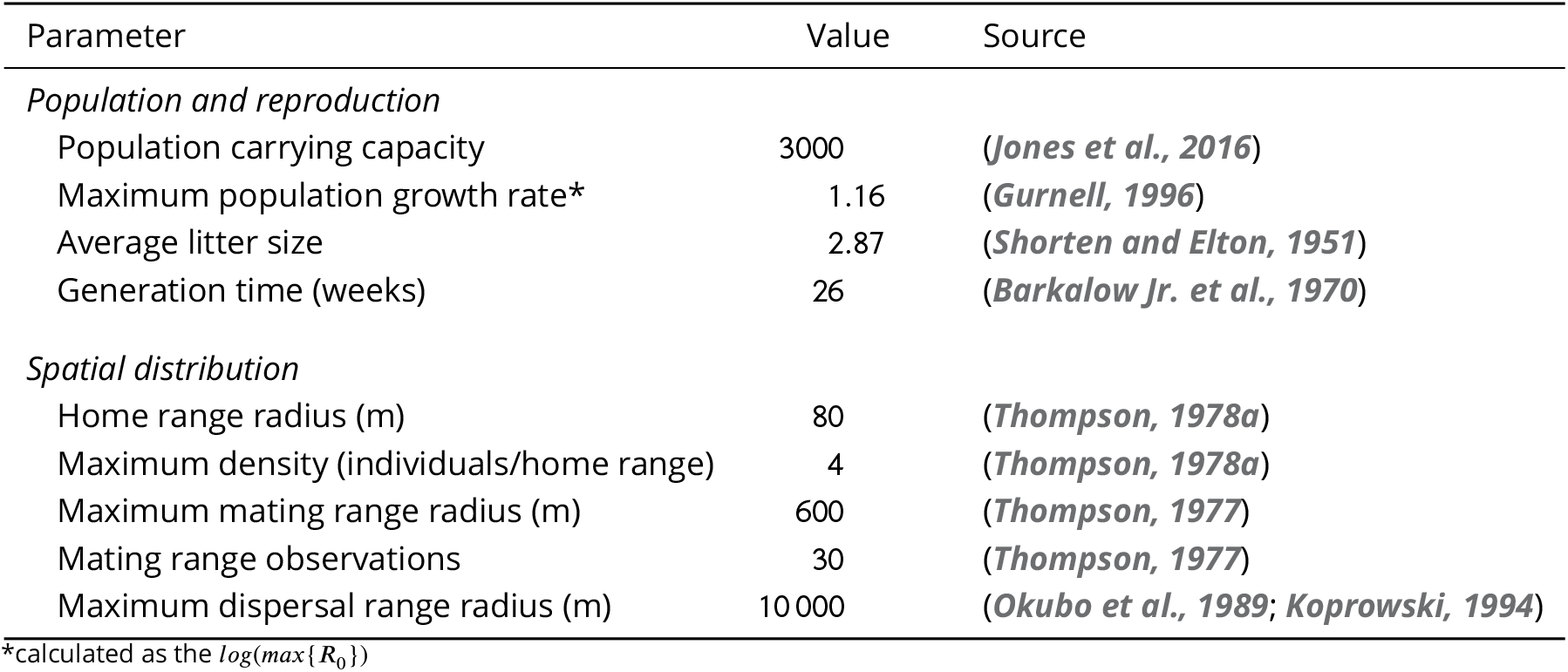
Key parameters used in the model for the grey squirrel. For the rest of the parameters, see the supplementary code.

| Parameter | Value | Source |
| --- | --- | --- |
| *Population and reproduction* |  |  |
| Population carrying capacity | 3000 | (*Jones et al., 2016*) |
| Maximum population growth rate* | 1.16 | (*Gurnell, 1996*) |
| Average litter size | 2.87 | (*Shorten and Elton, 1951*) |
| Generation time (weeks) | 26 | (*Barkalow Jr. et al., 1970*) |
| *Spatial distribution* |  |  |
| Home range radius (m) | 80 | (*Thompson, 1978a*) |
| Maximum density (individuals/home range) | 4 | (*Thompson, 1978a*) |
| Maximum mating range radius (m) | 600 | (*Thompson, 1977*) |
| Mating range observations | 30 | (*Thompson, 1977*) |
| Maximum dispersal range radius (m) | 10 000 | (*Okubo et al., 1989; Koprowski, 1994*) |

\*calculated as the $\log(\max\{R_0\})$

### Randomly mating model

For the randomly mating model, we added three additional features to the model of ***Prowse et al.*** (***2017***): cleave-and-rescue, daisyfield, and X-shredder. Cleave-and-rescue and daisyfield were not tested by ***Prowse et al.*** (***2017***), who only compared homing-based gene drives. We also modelled an X-shredder-cleave-and-rescue gene drive, but the homing-cleave-and-rescue was deemed more promising because the identification of a highly-specific spermatogenesis promoter remains a challenge. Also, X-shredder gene drives suffer from the formation of a population equilibrium instead of complete suppression. In addition to these three new features, we extended the supplementation functionality, beacuse daisyfield-based population suppression requires flexible supplementation.

#### 1. Cleave-and-rescue

In the model, we keep track of each gRNA-targeted site in cleave-and-rescue target genes and their functionality in each individual. The homing gene drive construct contains the recoded rescue copy of this target gene. All wildtype organisms start with two viable target genes, while gene drive organisms start with one viable target gene and one rescue. In general, after germline Cas-gRNA activity, viable target genes are cleaved and the rescue gene homes along with the gene drive. However, as with any sites targeted by a Cas-gRNA, it is possible that resistance alleles form after non-homolgous end joining and on occasion restore functionality of the target gene. Therefore, we implemented cleave-and-rescue gRNA multiplexing in the model. The probability that cleave-and-rescue target genes go from *i* to *j* functional cutting sites (*P*_*ij*_) is:

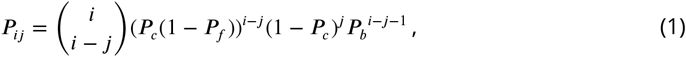

where *P*_*c*_ is the probability of cutting at a gRNA-targeted site, *P*_*f*_ is the probability of functional restoration in case of cutting, and *P*_*b*_ is the probability that a block of DNA in between two cutting sites is not removed. This formula consists of four factors: first, we multiply by all permutations of cutting sites, because their order is irrelevant. Second, we multiply by the probability that *i* − *j* cutting sites are all cut and repaired functionally. Third, we multiply by the probability that *j* sites remain uncut. Fourth, we multiply by the probability that no blocks of DNA in between cut sites were removed. We use *P*_*c*_ = 0.95 and *P*_*f*_ = 0.667 following ***Prowse et al.*** (***2017***). We estimated *P*_*b*_ from our unpublished data of 18 mouse embryonic stem cell lines, each cut simultaneously with Cas9 at two sites spaced 36 bp apart. In 3 out of 18 cases, the block of DNA in between the cut sites was not removed and therefore, we use a *P*_*b*_ of 0.2. All left-over probability (1− *P*_*ij*_) is the probability that a target gene is rendered non-functional. An organism needs to have exactly two copies of the target gene (recoded rescue or original) to be viable. We assumed that there is no embryonic Cas-gRNA activity. After random inheritance of parental alleles, we remove non-viable offspring.

#### 2. Daisyfield

We implemented daisyfield by tracking the number of daisyfield elements in the genome of each individual. Wildtype organisms start without any daisy elements and the number of daisy elements for gene drive organisms is a parameter in the model. Each daisy element contains both the homing and the cleave-and-rescue gRNAs and in case of gRNA multiplexing, it contains one of each different gRNA. Therefore, during germline Cas-gRNA expression, if no daisy elements are present, both homing and cleave-and-rescue can not occur. We assumed that daisy elements remain complete through every meiosis, so there is no crossing over in the middle of them. Also, we assumed that there is no linkage between daisy elements, that is, they are spaced far apart or located on different chromosomes. During inheritance, each daisy element from the parents has a 0.5 probability of being inherited to the offspring.

#### 3. X-shredder

Although the X-shredder is not a part of our final gene drive strategy, we implemented it in the model. The X-shredder gene drive is modelled on the Y-chromosome and skews the sex ratio of offspring towards males. The efficiency of this skew is a parameter in the model and is defined as the probability that offspring of a gene drive animal is male.

#### 4. Supplementation

We made two changes to the supplementation already implemented by ***Prowse et al.*** (***2017***). Instead of yearly suplementation of the same amount as the initial gene drive release, we added two parameters to vary supplementation amount and interval. Supplementation amount can be any percentage of the total population size, and supplementation interval can be any decimal number of years as long as they coincide with generations.

### Spatial modelling

For the spatial modelling, we added basic spatial functionality on top of the other additions to the randomly mating model of ***Prowse et al.*** (***2017***). We model a square, two-dimensional space and assume uniformly distributed resources such as food. The spatial functionality is comprised of four steps: spatial setup, distance-dependent mate allocation, offspring placement, and movement. The spatial setup is only done once at the start of the model and initiates everything necessary for spatial functionality. Mate allocation, offspring placement, and movement occur each generation, and their purpose is to reflect spatial life histories. Distance-dependent mate allocation ensures that squirrels who are close together are more likely to mate than squirrels further apart. Offspring placement demonstrates the location of birth and maternal care of individuals. Movement reflects the migration of individuals whenever overpopulation occurs in an area. With several parameters shown in ***Table 1***, this spatial functionality can be adapted to reflect the spatial life history of many species. Additionally, we have added spatial placement strategies for supplementation.

#### 1. Spatial setup

The first step in spatial modelling is to determine the size of the area in which the simulations take place. As we use a square two-dimensional space, we need to know the length of the side of this area *A*. We calculate *A* using the carrying capacity of the population *K*, the radius of the home range of the organism *r*, and the density at carrying capacity *D*:

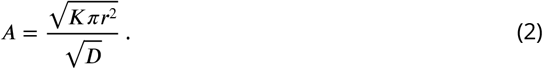

Essentially, this formula transforms a circular home range radius into an area, multiplies it by the number of individuals, transforms it into the length of a square area, and makes it smaller according to the density at carrying capacity. Using this formula, the area is exactly large enough to hold *K* number of individuals at *D* density. In this two-dimensional area, we track the *x* and *y* coordinates of individuals. Each individual starts at a random location within the area. Where gene drive individuals are placed depends on the placement strategy.

#### 2. Distance-dependent mate allocation

During the reproduction step of the model, instead of random mate allocation, we use distance-dependent mate allocation. We do this in three steps. First, we calculate the Euclidian distance between all females and males. Second, we use a Gaussian radial basis function to calculate the probability of a male approaching the female to mate (*P*_*a*_), depending on the distance *s* between them:

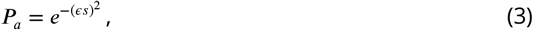

where the value *ϵ* determines the shape of the radial basis function and is calculated from the mating range parameter. In the case of the grey squirrel, the maximum observed mating range was 600 out of 30 observations (***Thompson, 1977***). Therefore, we assumed that the probability of a mating range of 600 was 1/30 and from this, we calculate *ϵ*. Third, from the males that do approach the female, we choose a random one as the father of the offspring. In the case that no males approach the female, she doesn’t reproduce.

#### 3. Offspring placement

We place offspring at the location of the female at the moment of reproduction.

#### 4. Movement

In grey squirrels, migration is the driving force behind a stable population size (***Thompson, 1978b***). Therefore, we implemented density-dependent migration and not density-dependent mortality. In the model, we make a distinction between the movement of migrants and residents. Firstly, we determine which individuals migrate and which remain as residents. This distinction is density dependent, that is, the density at the location of an individual determines the probability that they migrate (*P*_*m*_):

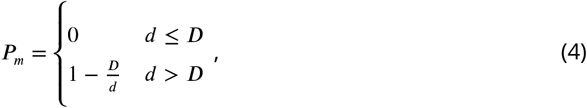

where the local density *d* and the density at carrying capacity *D* are measures of the number of individuals that are in the home range of an individual. Therefore, when the local density is below maximum density, individuals will not migrate. When the local density is higher than the maximum density, the probability of migration is equal to the proportion of individuals that need to migrate to leave the local density at the maximum density. Next, for both the resident and the migrant movement, we choose a direction and a distance to determine a new location. We choose a random direction and a distance from two seperate gamma distributions for residents and emmigrants with shape and scale parameters: *distance ~ Γ*(*k, θ*) ≡ Gamma(5*, r*/5) for residents and *distance ~ Γ*(*k, θ*) ≡ Gamma(5, 3*r*/5) for migrants, *r* being the home range. We use a broader distribution for migrants than for residents as migrants tend to travel greater distances (***Thompson, 1977***). The residents move to a random location in a single step. If the new location is out of the boundaries of the spatial space, we pick a new direction and distance. In contrast, migrants move in multiple steps within a certain migrational range to a place where there is space available, that is, where the local density *d* is lower than the density at carrying capacity *D*. The migrant searches for a new location in a lazy manner, which means that an animal will first try nearby locations, and incrementally migrate further if necessary. In each step, we pick a random direction and add a new distance from the gamma distribution to the previous distance. If the maximum migration distance is surpassed, the distance is set to zero and the process starts again. To ease the computational burden of this algorithm, we limit the number of steps to 50 and then, we keep the last location regardless of density.

#### 5. Supplementation

The placement of individuals for supplementation is important. Therefore, we have implemented five placement strategies that can be used, although further exploration of this aspect is interesting. The six placement strategies are: middle of the area, mean of population location, mode of population location, random location at each supplementation, divided into 10 groups and placed at random locations at each supplementation, and divided into 10 groups and placed as a moving front in 10 steps.

## Acknowledgments

We thank Craig Shuttleworth for all his squirrel-related expertise and advice.

## Data accessibility

Code and data are available from the Highlanderlab gitlab: https://git.ecdf.ed.ac.uk/HighlanderLab_public/nfaber_squirrel_gd.

---

**Figure 3–Figure supplement 1.**
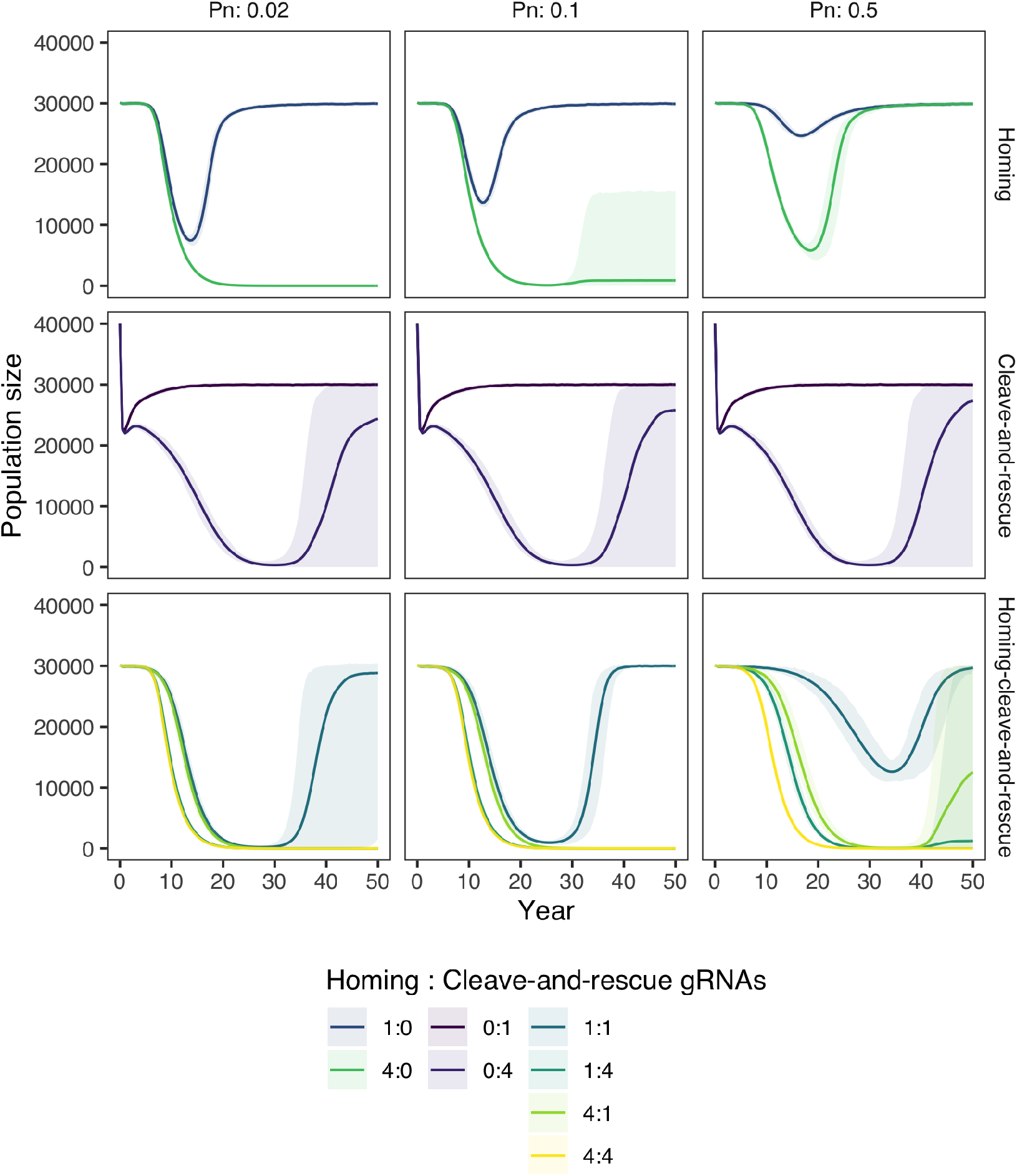
The same as **Figure 3**, but run in a big population with a carrying capacity of 30,000 instead of 3,000. Population size over time after the introduction of gene drive squirrels with either a standard homing, a standard cleave-and-rescue, or a homing-cleave-and-rescue gene drive to a population with carrying capacity 30,000. All simulations are based on a single release of 100 squirrels is done, other than the standard cleave-and-rescue gene drive, which requires a release of 10,000 squirrels. Lines represent the average population size over 100 model replications, while opaque ribbons represent the 95% quantiles. The model was run with 3 different rates of NHEJ repair during homing (*Pn*) and with different numbers of gRNAs for the homing and the cleave-and-rescue components of the gene drive.

**Figure 3–Figure supplement 2.**
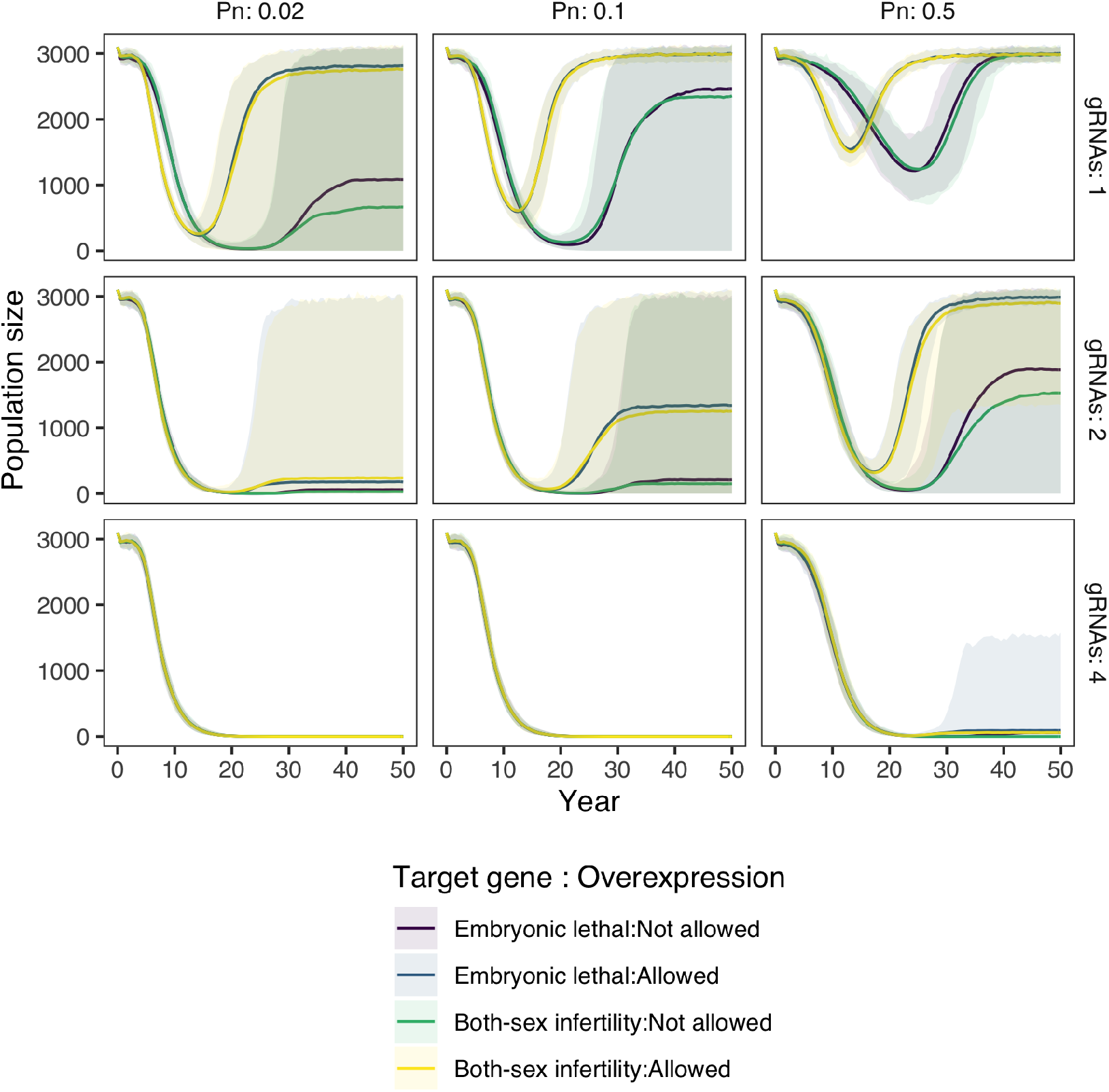
An exploration of which type of gene is best targetted by the cleave-and-rescue part of the gene drive: both-sex infertility or developmental non-viability, and overexpression biologically tolerable or not. Parameters are kept the same as in **Figure 3**, except that we used 1 gRNA for the homing part of the gene drive, and either 1, 2 or 4 gRNAs for the cleave-and-rescue part. Population size over time after the introduction of 100 gene drive squirrels with a homing-cleave-and-rescue gene drive to a population with carrying capacity 3,000. Lines represent the average population size over 100 model replications, while opaque ribbons represent the 95% quantiles. The model was run with 3 different rates of NHEJ repair during homing (*Pn*).

**Figure 4–Figure supplement 1.**
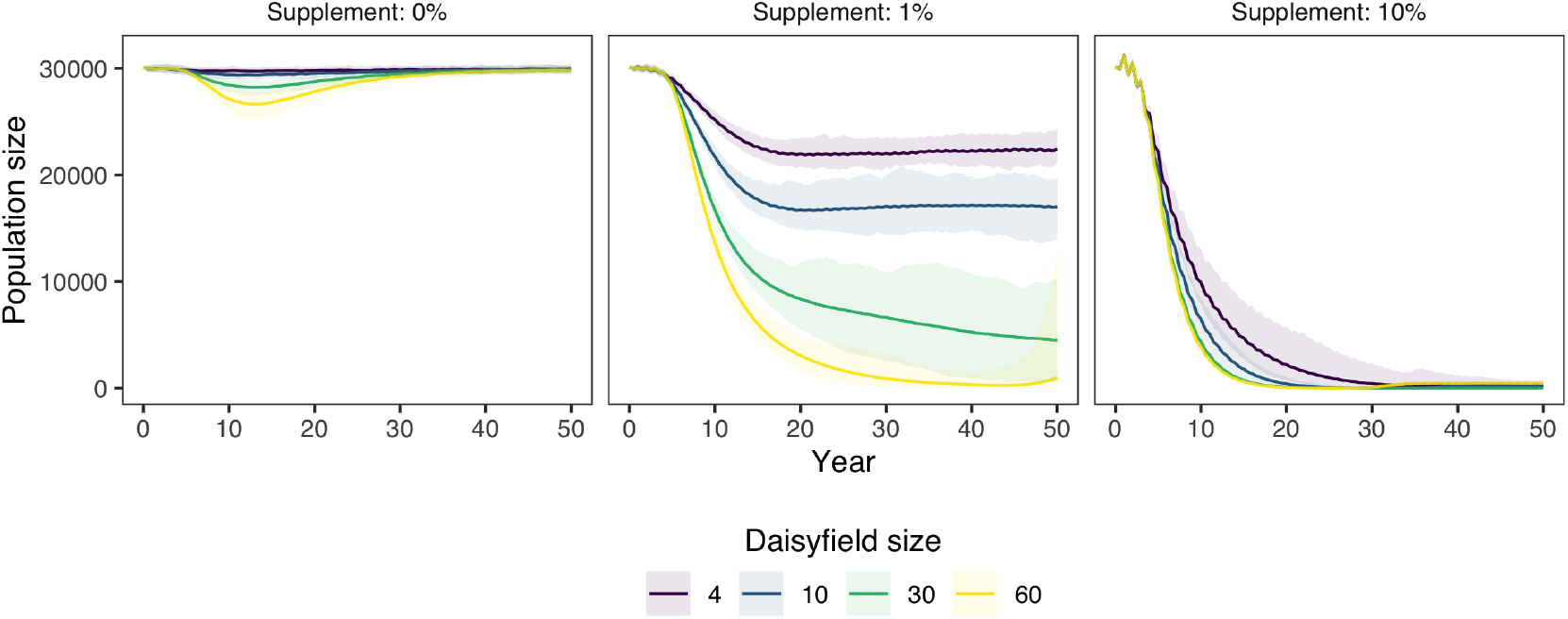
The same as **Figure 4**, but run in a big population with a carrying capacity of 30,000. Population size over time after the introduction of 100 squirrels with a HD-ClvR gene drive. The model was run with an NHEJ rate (*Pn*) of 0.02, 1 homing gRNA, and 4 cleave-and-rescue gRNAs. Gene drive squirrel supplementation was done yearly, the amount being a percentage (0, 1, or 10%) of the total population size at that moment. Lines represent the average population size over 100 model replications, while opaque ribbons represent the 95% quantiles.

**Figure 4–Figure supplement 2.**
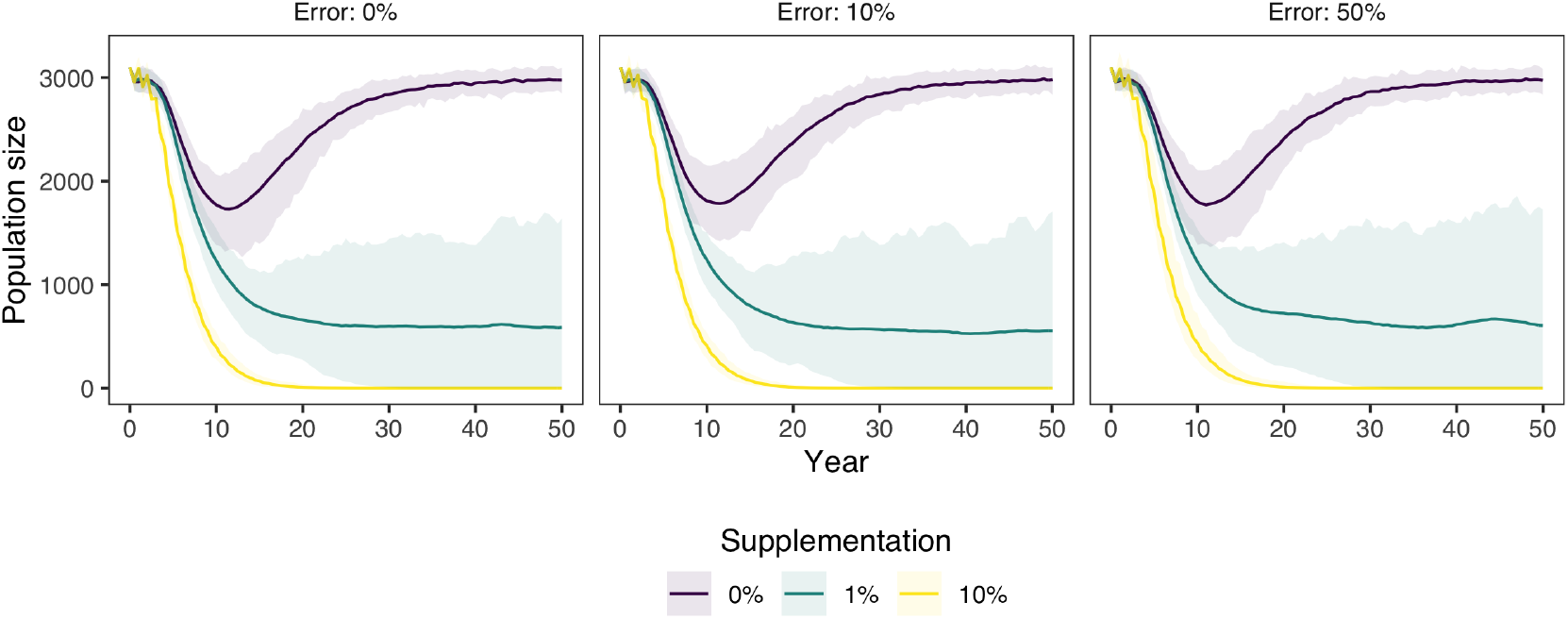
The same as **Figure 4**, but instead of an accurate estimate of the population size for supplementation, a certain level of error is introduced. The error is defined on a yearly basis as a normal distribution with the true population size as mean and a certain percentage of the true population size as standard deviation. Population size over time after the introduction of 100 squirrels with a HD-ClvR gene drive to a population of carrying capacity 3,000. The model was run with an NHEJ rate (*Pn*) of 0.02, 1 homing gRNA, and 4 cleave-and-rescue gRNAs. Gene drive squirrel supplementation was done yearly, the amount being a percentage (0, 1, or 10%) of the total population size at that moment, plus the abovementioned error. Lines represent the average population size over 100 model replications, while opaque ribbons represent the 95% quantiles.

**Figure 4–Figure supplement 3.**
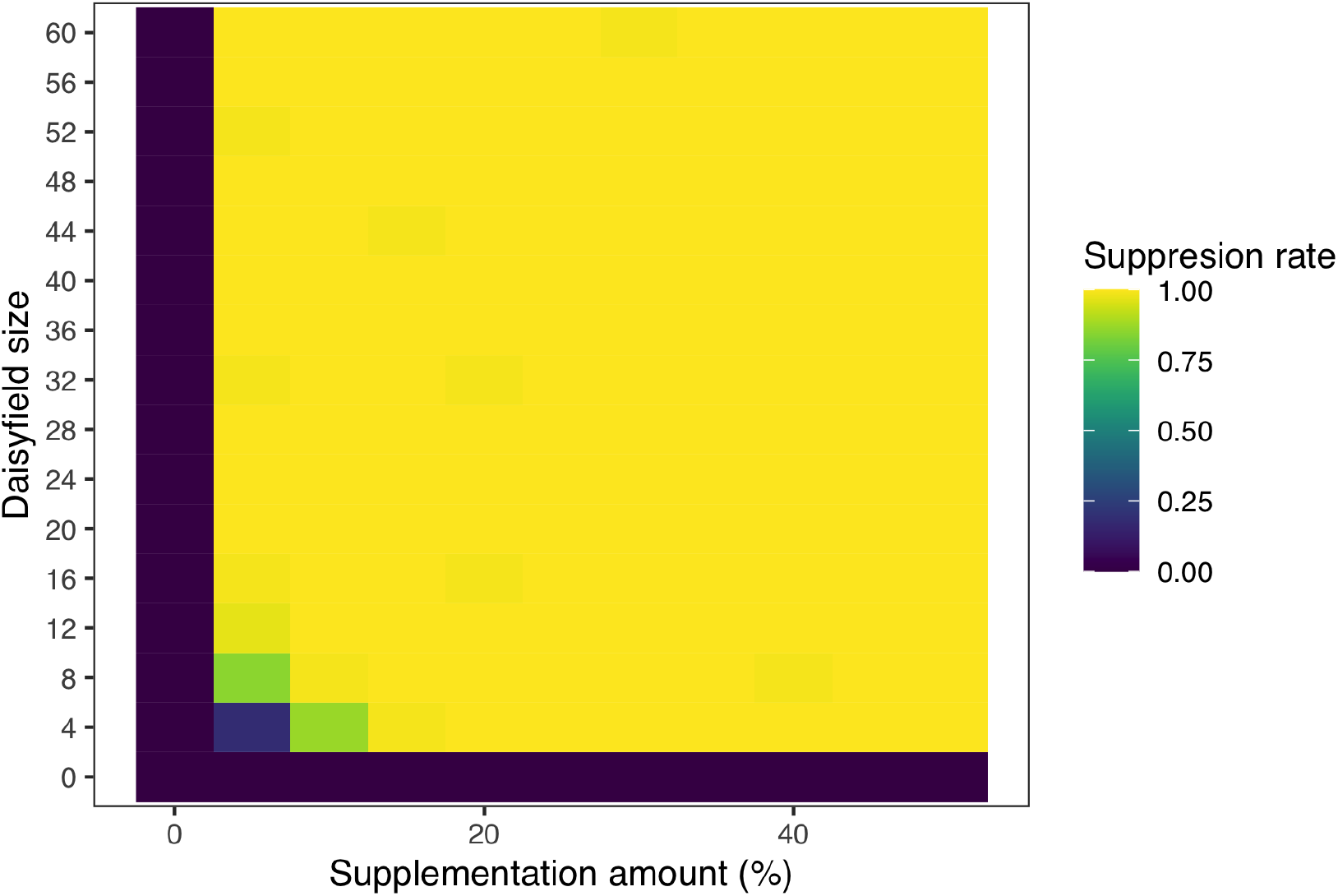
The same as **Figure 4**, but ran with a range of supplementation amounts and daisyfield sizes. Suppression rate is defined as the proportion of populations (out of the 100 repetitions of the model) that were completely suppressed after 50 years. Suppression rate after the introduction of 100 squirrels with a HD-ClvR gene drive to a population of carrying capacity 3,000. The model was run with an NHEJ rate (*Pn*) of 0.02, 1 homing gRNA, and 4 cleave-and-rescue gRNAs.

**Figure 5–Figure supplement 1.**
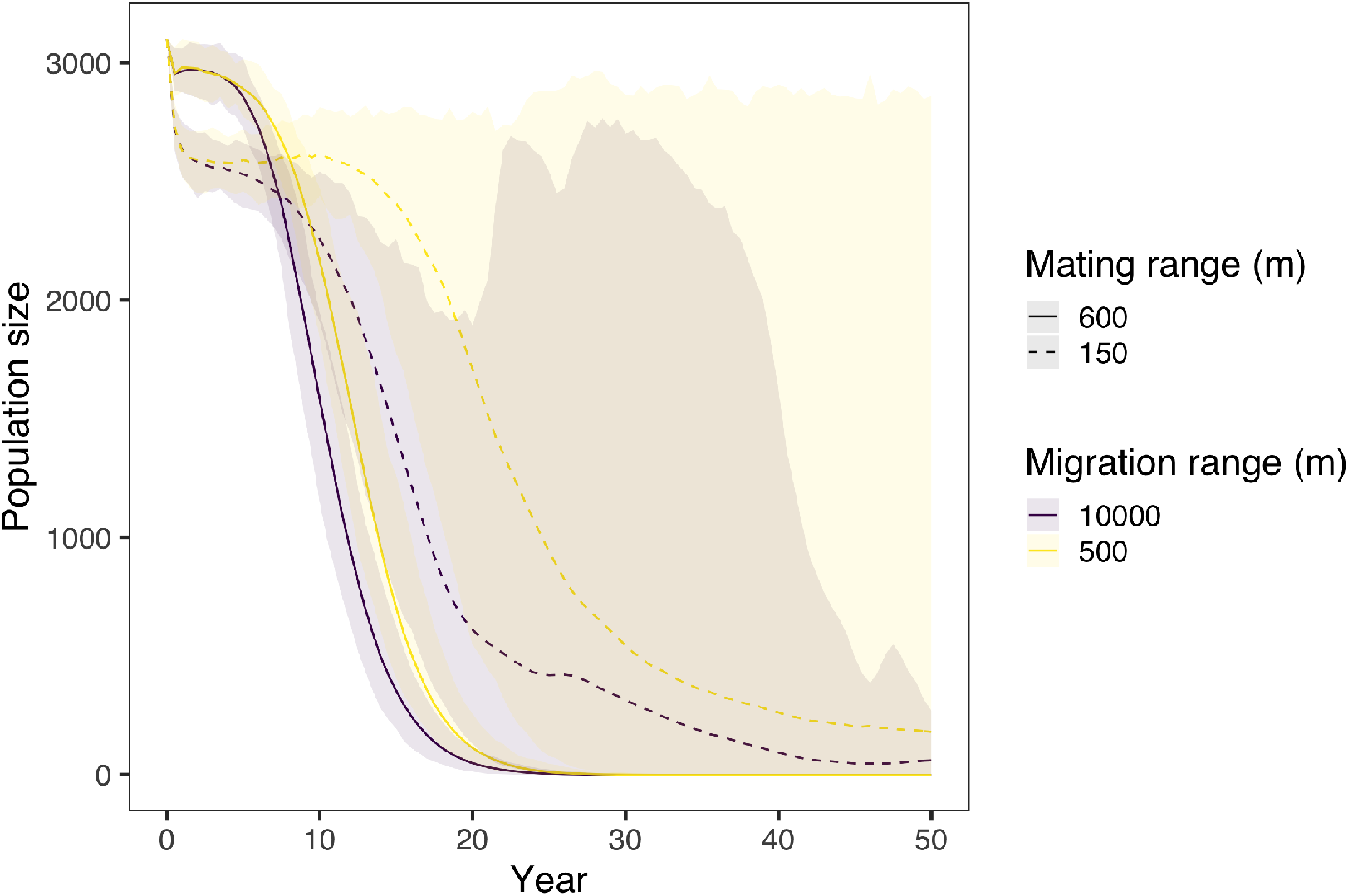
A sensitivity analysis of two crucial parameters in our spatial model (**Figure 5**): mating range and migration range. We model population size over time after the introduction of 100 squirrels with a homing-cleave-and-rescue gene drive with 1 homing gRNA and 4 cleave-and-rescue gRNAs. An NHEJ rate (*Pn*) of 0.02 was used. In the spatial model, gene drive squirrels were placed in the middle of the area. Lines represent the average population size over 100 model replications, while opaque ribbons represent the 95% quantiles.

